# Automated bird flight pattern extraction and classification using machine learning

**DOI:** 10.64898/2026.03.17.712367

**Authors:** Milica Ostojic, Sarab Sethi

## Abstract

With bird populations across the world being impacted by ever-growing anthropogenic pressures, reliable monitoring is essential to help halt or reverse declines. Existing visual bird monitoring approaches, which employ cameras or radars to deliver automated and large-scale monitoring data, face a variety of issues. Image-based species classification is only possible if the fine-scale features of a bird are clear, which can be difficult to achieve in real monitoring contexts without expensive, high-resolution cameras due to occlusion and lighting. Radar and video-based approaches which analyse longer-term flight behaviour over the course of seconds can achieve more reliable results in real monitoring contexts, particularly from greater distances, but still require expensive equipment and do not account for all the possible types of flight patterns. Here we present a novel approach to track a wide range of bird flight patterns using inexpensive equipment. As a proof-of-concept, we demonstrate how our approach can be used to classify birds between four species, Red Kite, Kestrel, Black-Headed Gull and Sparrowhawk, which represent four different types of flight patterns. The balanced accuracy of the classification is 0.5583, with a recall and precision per species that range from 0.2640-0.7750 and 0.4583-0.5962, respectively. Our proof-of-concept study demonstrates how new and existing visual bird monitoring systems can leverage flight patterns to deliver species-level insights at lower costs and on larger scales than before.

## 1. Introduction

Bird population sizes are decreasing globally, with approximately 48% of global bird species currently suspected of being in decline (1), driven by habitat loss, land use change and degradation (2), as well as other anthropogenic pressures such as climate change (3). This global trend has manifested as a net loss since 1970 of almost 3 billion birds in North America (1), and 73 million birds in the UK (a decrease of almost a third) (4). Wild birds contribute many ecosystem services, including seed-dispersion, pollination, ecosystem engineering and linking of global ecosystem processes due to the migration of certain bird species (1). They are also important bioindicators, which are organisms that can be used to monitor the health of an environment, such as whether an area has been impacted by pollution or contamination (5).

Reliable monitoring of birds is thereby becoming ever more imperative as pressure on global ecosystems increases. While monitoring efforts are slowly scaling up, there are still significant spatial (6) and temporal (7) data gaps. Filling these data gaps is essential to drive effective policy and conservation work, as is monitoring the impact of interventions, such as policy changes (8) or rewilding efforts (9). To do this, a change in approach is needed. Traditional monitoring techniques, such as tagging (10), manual point counts and transects (11), can be considered invasive, as well as labour and resource intensive (11). Alternative, automated methods are therefore gaining popularity due to their ability to increase the scale, cost-effectiveness and time-efficiency of monitoring, allowing scientists and land managers to monitor birds and therefore environmental health across greater spatial and temporal scales with improved accuracy.

Existing automated methods of monitoring birds include acoustics and visual systems, both of which have their own advantages and shortcomings. Acoustic methods improve the scalability of bird monitoring, both spatially and temporally (12), and decrease disturbance (12) compared to traditional methods like point counts. However, acoustic monitoring approaches struggle to identify silent species, species vocalising in noisy environments, and individuals which are far from the listening devices (13). Automated visual identification, using data from satellite imagery (14), drones (15), and camera traps (16) can provide a complementary and similarly scalable approach to bird monitoring with minimal disturbance. Most existing visual bird monitoring systems use machine learning models to predict bird species from their appearance in static snapshots (17). However, image classification requires high resolution, close-up images to capture the fine details in the appearance of a bird. These images are difficult to obtain in real-world monitoring applications due to the unpredictability of a bird’s orientation, occlusions from the environment, and the cost of high quality cameras and lenses (18)).

Approaches that go beyond snapshots, and take into account some aspects of bird flight to provide species monitoring data use radar (19,20), video (16) or both in combination (22). In radar systems, detection can be achieved by combining signal strength and flight speed to determine whether the detected object is a bird and to estimate its size (19). Detection can also be achieved using the temporal pattern of the echo signature (19,20). The radar signal changes depending on the shape of the bird (19), which varies during flapping motion as the bird flaps its wings. The distinctive flapping pattern produced can be used to identify a bird over other flying objects (19,20) and differentiate between groups of birds, for example passerines and waders (20). Video-based systems can take a similar approach to radar. For example, in a study conducted by Zhang *et al* (2008) (21), the flapping pattern is extracted by using the size of the bird in the video field to determine at which point in the flapping period the bird is – when the wings are flat, the size is at its largest, whereas as the bird moves its wings up and down, its size in the frame varies. Visual and radar data can also be combined to achieve species identification, like in the system developed by Niemi *et al* (2018) (22).

Radar is used to detect incoming birds and to provide the distance to the bird, its velocity and its trajectory. A camera with a telephoto lens and a motorised video head is then used to capture a series of images of the bird, which are classified by species using a CNN and segmented to estimate the size of the bird in combination with the radar parameter of the distance to the bird. All this information is then combined with the velocity of the bird to determine a final prediction.

However, there are significant drawbacks to radar and video-based systems. Radar data includes signals from many other flying objects, such as insects, bats and even weather phenomena (19), some of which produce similar signatures to birds (19). Furthermore, by only using flapping patterns to differentiate birds from other flying objects or to identify species (19,20), radar systems can overlook birds that exhibit gliding behaviour (19). Larger birds often glide for long periods without flapping (23), and therefore a flapping pattern cannot be extracted even though such birds produce stronger radar signals than smaller and mid-sized birds (19). The existing video-based method developed by Zhang *et al* (2008) (21) could take into account gliding and mitigate the issue with radar, but faces a separate significant limitation in that the camera has to maintain a consistent relative angle to the bird within a small range. Combining radar and visual hardware may compensate for some of the issues with the separate systems but does still require expensive and high-performance visual hardware, such as the telephoto lens camera used by Niemi *et al* (2018) (22).

In this study, we develop a novel approach that considers the mechanics of bird flight to analyse both flapping and gliding motion (thereby including larger birds in its classification) with the use of inexpensive equipment and minimal limitations on the video input. The system we present is a proof-of-concept prototype capable of detecting a bird within a video of a single bird in flight and classifying between four species that represent different types of flight pattern. We achieve this by processing each video using a combination of custom and off-the-shelf machine learning models to generate the flight pattern of the bird from the individual flaps of its wings, and then we extract the features needed to identify its species.

## 2. Materials and Methods

### 2.1. Overview of approach

Below is an overview of our approach (Figure 1). In this methods section, we will give more details on each step shown.

**Figure 1:**
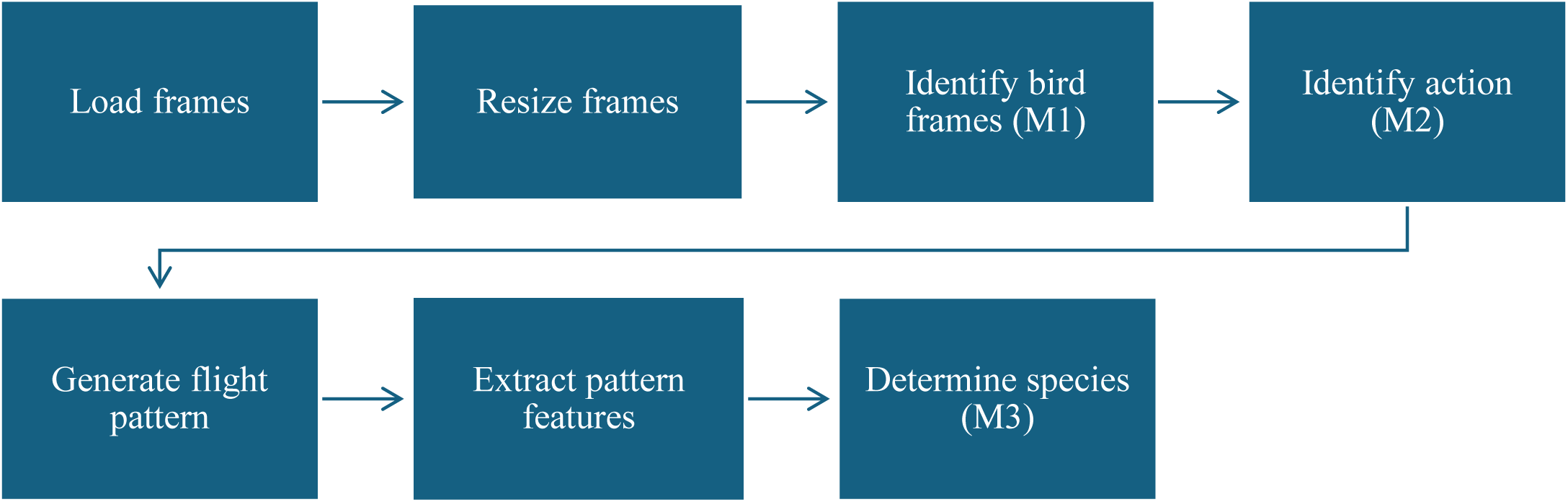
Overall pipeline for identifying species from a video input of a single flying bird.

### 2.2. Upstroke vs downstroke

During flapping motion, a bird will perform a periodic pattern of upstrokes and downstrokes (24). A downstroke is when the wings are moving downwards, and they are outstretched to produce lift and propulsion (24). An upstroke is when the wings are moving upwards and being drawn in towards the body (24), thereby bending at the wing joint (24) and reducing drag to preserve the lift and thrust produced by the downstroke (24). During gliding motion, a bird will tend to keep its wings in either an upstroke position or a downstroke position for a prolonged period. The choice may depend on environmental factors like wind strength. Gliding in downstroke allows the bird to generate more lift as the wings have a greater surface area in downstroke (25). However, in stronger headwinds, birds will partially retract their wings into an upstroke position to optimise lift to drag ratio by reducing their surface area to decrease drag while still maintaining some lift (26). In our approach, we use the building blocks of individual upstrokes and downstrokes to generate the flight pattern of a bird.

### 2.3. Glossary

- M1 – bird detection model
- M2 – action classification model (upstroke or downstroke)
- M3 – species identification model (Red Kite, Kestrel, Black-Headed Gull, or Sparrowhawk)

### 2.4. Datasets

We assembled the below four datasets (and one subset) for this study by drawing upon a variety of public datasets.

The action classification model (M2) training dataset was formed from 8,699 images of birds in flight manually selected from the North American Birds dataset (NABirds) (27,28) and the 2019 iNaturalist dataset (29). The NABirds dataset, comprising of 48,562 images (28), was created by Van Horn *et al* as a collaboration between Cornell Lab of Ornithology, Cornell Tech, Caltech, Birgham Young University and citizen scientists (27). Experts within the birding community curated the dataset from images contributed by members of the community (28) via the All About Birds website (27). The 2019 iNaturalist dataset, comprising of 268,243 images (29), was collated from images contributed by citizen scientists via the iNaturalist website and app (30). We manually annotated each bird within the selected images from these datasets (some images had more than one bird) as being either in upstroke or downstroke to produce a new labelled dataset. There were 5,754 downstrokes and 4,421 upstrokes.

The M2 full-frame test dataset, in which the bird in each image takes up most of the frame, comprises of 639 images of birds in flight manually selected from the CUB-200-2011 dataset by Wah *et al* at Caltech (31). The CUB-200-2011 dataset was created by Wah *et al* by performing a search on Flickr and then having users of Mechanical Turk (Amazon’s crowdsourcing website for hiring remote workers for on-demand tasks) filter the results (31). We labelled the selected images with upstroke and downstroke. There are 352 downstrokes and 287 upstrokes.

A 5-second video clip dataset was used for Group Shuffle Split cross-validation of the species identification model (M3). It was created by first generating 561 5-second clips (297 Red Kite, 160 Kestrel, 64 Black-Headed Gull, 40 Sparrowhawk) from 80 videos of birds in flight (40 Red Kite, 23 Kestrel, 10 Black-Headed Gull, 7 Sparrowhawk) contributed by members of the birding community via the Macaulay Library at the Cornell Lab of Ornithology (32). The 5-second clips from each video were generated by stepping a window of 5 seconds through each video with a step size of 2 seconds, so the resulting clips overlap. 40 clips per species were then randomly selected (with each clip originating from a different longer video) to create the balanced 5-second video clip dataset for cross-validation of species identification.

A test subset of the clip dataset (bird vs non subset) was used to extract frames for testing the bird detection model (M1) and the action classification model (M2). To ensure that there were empty background frames to test with, the clips in the balanced 5-second clips dataset that had periods in which the bird was off-screen were manually selected. If there were less than 4 clips containing frames without a bird (non-frames) for each species (and originating from different longer videos), additional clips were randomly selected to add up to 4. If there were more than 4 clips with non-frames for a species, 4 clips (originating from different longer videos) were randomly selected from those with non-frames. All the frames, showing either a bird or an empty background, from the 16 resulting clips were then extracted and labelled. Non labels were assigned for frames that do not contain a bird, which in this dataset were empty background frames. The dataset labelled with bird vs non was used to test the ability of M1 and M2 in classifying frames with a bird and empty background frames. The dataset labelled with upstroke vs downstroke vs non was used to test the ability of M2 to classify between upstrokes, downstrokes and empty background frames.

A dataset of other flying objects was assembled from 538 images of single aeroplanes, helicopters and drones manually selected from the AOD 4 Dataset for Air Borne Object Detection created by Soni *et al* (33,34). The AOD 4 Dataset was collated from multiple sources (YouTube-8 M, Anti-UAV and a dataset by Ahmed Mohsen hosted on Roboflow) with all the 22,516 images uniformly resized to a resolution of 1024 x 1024 (33). All the images selected from the AOD 4 dataset were labelled with non as none of them contain a bird, which is consistent with the definition of a non-label in the above subset of the 5-second clips dataset. The resulting dataset of other flying objects was therefore able to be combined with the frames labelled bird vs non from the subset of the 5-second clips dataset to test M1 and M2.

### 2.5. Bird detection and motion classification

M1, the model used to identify the frames with a bird present, is a pre-trained object detection model available on the PyTorch Hub for researchers (Single Shot MultiBox Detector model for object detection (35)), which outputs the class and bounding box of an object in the input frame (35). It is based on a ResNet-50 model (35) and has been trained on the COCO (Common Object in Context) dataset (36), which is a large-scale image dataset used widely for training object detection models. To convert the multiclass model into a binary model, only “bird” predictions are considered when comparing the confidence of a prediction to the minimum threshold, regardless of whether there are predictions for other classes with higher confidences, leading to only two options – “bird” and “non”.

The input size of the model is 300x300 pixels (35) so, once the individual frames of the input video have been loaded, they are rescaled to match this format. Preliminary tests suggested that padding with white or black pixels to retain the same aspect ratio produced slightly worse performance than rescaling without padding.

To compensate for the false negatives from M1, we explored using a TLD (tracking, learning and detection) tracker from the Python OpenCV 4.8.1 library. The tracker was re-initialised with a bounding box from M1 for each frame that was predicted as a bird by M1. The tracking algorithm could then track the bird between these frames. However, we found that including object tracking resulted in an ROC that didn’t change with increasing threshold from right to left like a typical ROC (see Appendix A for Figure A.1). This suggests that the performance of object tracking depends significantly on the specific combination of frames within a video that M1 identifies as “bird” and therefore the performance is unpredictable for new data. Furthermore, for the chosen M1 threshold, the addition of object tracking didn’t significantly change the performance of the bird detection stage. Therefore, we opted to use M1 without object tracking for bird detection.

M2 was trained using images in which the bird takes up most of the frame, due to the available training datasets. Therefore, to match the training data, we trialled cropping the frames with a detected bird around the bounding box determined by M1 before inputting into M2, but the performance reduced rather than improved, so instead we feed the original frames to M2.

Once the frames with a bird have been identified, M2 determines the action of the bird for each frame in which the bird is present. We used the Python detecto (37) module’s transfer learning implementation to train a custom object detection model with two output classes – upstroke or downstroke – built on the ResNet-50 architecture (the original model was pre-trained on the COCO dataset). It is possible that both an upstroke and a downstroke can be detected by the model within an input frame, in which case, we take the label with the highest confidence. If the model is unable to find any upstrokes or downstrokes, we assign the frame a “non” label.

### 2.6. Flight pattern characterisation and species classification

A time series of “upstroke”, “downstroke” and “non” labels is generated, forming the flight pattern for the full video. If there are more than 5 consecutive “non” labels, which is equivalent to 0.2 seconds for a 25fps video, it is assumed that the bird is off-screen and therefore the flight pattern is split. The rest of the “non” labels are then removed from each resulting pattern, which does not significantly affect the periodicity of the upstrokes and downstrokes or the timing of the flight modes (flapping and gliding), given that the removed sections are less than 0.2 seconds in duration. Any resulting flight patterns that are less than 2 seconds long are discarded for the downstream species prediction task.

Features of each flight pattern generated from the input video are then extracted by a deterministic algorithm. To identify the flapping and gliding sections, the algorithm first determines all the points at which the input pattern switches between upstroke and downstroke, known as switching points. The average of all the time differences between switching points is then calculated as an estimation of half the flapping period (FP_1/2_), where one flapping period is the time taken for one up-and-down movement of the wings. The value of FP_1/2_ is then used to determine which switching points are part of a flapping section and which mark the start or end of a gliding section. Each switching point is within a flapping section if the time until the next switching point (Δt_SP_) fulfils the criteria: Δt_SP_ < min(5(FP_1/2_), 1 sec), otherwise the switching point marks the end of a flapping section and the start of a gliding section. From this, for each flight pattern we calculate the following features: average flapping time, average gliding time, average flapping upstroke time, average flapping downstroke time, flapping to gliding ratio, and gliding to flapping ratio. Any features that are calculated as “nan” values (for example, due to dividing by zero), are reassigned to a value of 0.

The final model, M3, is a Random Forest Classifier that determines the species from the features of a flight pattern. Species classification is currently limited to four species – Red Kite, Kestrel, Sparrowhawk and Black-Headed Gull. M3 was tested using Group Shuffle Split cross validation (10 splits, 0.8/0.2 train-test split) of the 5-second clips dataset. The clips have been split into groups based on the original video they were generated from, to ensure no overlap in video content between the train and test datasets.

## 3. Results

We first conducted tests to determine whether an additional model (M1) was required for bird detection on top of the custom action classification model (M2). When differentiating between frames with a bird and empty background frames from the 5-second clips test subset (Figure 2a), the action classification model (M2) performed slightly better than the bird detection model (M1), with an AUC of 0.9577 compared to an AUC of 0.9404 for M1. However, M2 performed far worse than M1 when differentiating between birds and other flying objects (Figure 2b), with an AUC of only 0.7078 compared to an AUC of 0.9069 for M1.

**Figure 2:**
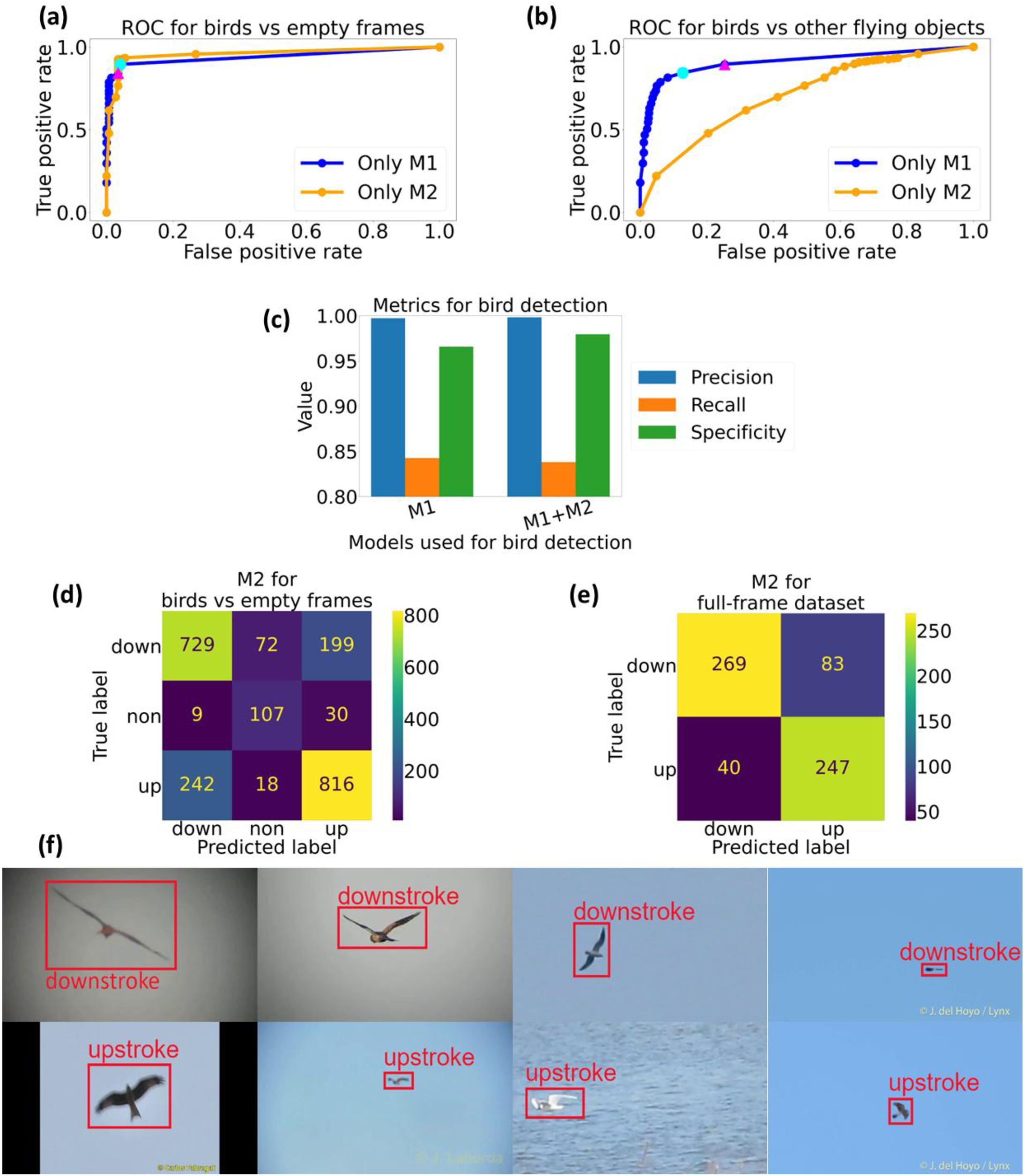
Bird detection and subsequent motion classification. (a) Receiver Operating Characteristic (ROC) for 5-second clips test subset, classifying birds vs empty frames, for M1 only (blue) and M2 only (orange) with the optimal threshold for M1 highlighted by a cyan circle (0.05). The optimal threshold for M1 when classifying birds vs other flying objects (0.1) is highlighted by a magenta triangle. Using a threshold of 0.1 for M1 slightly decreases true positive rate. (b) ROC for 5-second clips test subset combined with the other flying objects dataset, classifying birds vs other flying objects, for M1 only (blue) and M2 only (orange). M1 has an AUC of 0.9069 whereas M2 has an AUC of 0.7078. M2 is unable to reliably classify birds vs other flying objects, but M1 can at an optimal threshold of 0.1 (cyan circle). Using the optimal threshold for birds vs empty frames of 0.05 (magenta triangle) would significantly increase false positive rate compared to using a threshold of 0.1. (c) Bar chart showing precision, recall and specificity of each stage of bird detection. For solely M1 at a threshold of 0.1, precision is 0.9971, recall is 0.8425, and specificity is 0.9658. For M1 + M2, precision is 0.9983, recall is 0.8382 and specificity is 0.9795. M2 is slightly detrimental to the overall recall of bird detection but improves the specificity. (d) Confusion matrix for M2 on 5-second clips test subset. An accuracy of 0.7435 and a weighted precision of 0.7486. (e) Confusion matrix for M2 on full-frame dataset. An accuracy of 0.8075 and a weighted precision of 0.8157. (f) Examples of downstrokes and upstrokes identified by M2 for each species (left to right: Red Kite, Kestrel, Black-Headed Gull, Sparrowhawk) on frames from the 5-second clips test subset.

Therefore, we use M1 and M2 in combination. The optimal threshold for M1 when differentiating between birds and other flying objects is 0.1 (cyan circle on the blue line in Figure 2b), whereas for differentiating between birds and empty background frames, the optimal threshold is 0.05 (cycle circle on the blue line in Figure 2a). Given that there is a fairly significant increase in false positive rate when using a threshold of 0.05 for classifying birds and other flying objects (highlighted by the magenta triangle in Figure 2b), but only a slight decrease in true positive rate when using a threshold of 0.1 for classifying birds and empty background frames (highlighted by the magenta triangle in Figure 2a), M1 is used at a threshold of 0.1.

Our next test was to determine the overall performance of bird detection with both M1 and M2 used in combination. Figure 2c demonstrates how the addition of M2 to M1 results in a small drop in recall but an increase in specificity. The drop in recall is due to M2 occasionally being unable to detect an upstroke or a downstroke within a frame that M1 has correctly identified as having a bird, leading to a “non” label being assigned even when a bird is present. However, the increase in specificity indicates that M2 reduces the number of false positives, which also have a significant impact on downstream analyses.

Finally, we determined the performance of the action classification stage. M2 had an accuracy of 0.7435 and a weighted precision of 0.7486 for differentiating between upstrokes, downstrokes and non when tested on individual frames from the 5-second clips test subset (Figure 2d, 2f). When tested on the full-frame dataset instead, with images that were more similar to the M2 training dataset, M2 had a higher accuracy of 0.8075 and higher weighted precision of 0.8157 (Figure 2e).

For all of the bird detection results, it should be noted that since we are using object detection models, occasionally M1 and M2 outputted a “bird” class for a frame that had a bird in it, without correctly identifying the bounding box of the bird in the frame. For M1, this is a non-issue as all frames assigned as “bird” will be passed to M2 regardless of the location of the predicted bounding box. For M2, this may have an impact on the results as an incorrectly placed bounding box means that the model has not identified the bird and therefore the flight action (upstroke/downstroke) correctly, leading to errors in the flight pattern generated downstream. However, this appears to be a very minor issue. When generating the M2 ROC, the bounding boxes were manually checked for all the test frames, and only 1.8% had this issue.

Figures 3a-d show an example of a generated time series flight pattern for each species, alongside a series of frames from the clips the patterns were generated from. In each pattern, the flapping sections (pink) and switching points (red stars) are highlighted. The confusion matrix for Group Shuffle Split cross-validation of M3 on the 5-second clips balanced dataset is shown in Figure 3e, and the metrics calculated per species from the confusion matrix is shown in Table 1. The Red Kite had the best performance overall with the highest F1 score (0.6739). The Sparrowhawk had the lowest performance overall with the lowest F1 score (0.3350). As is clear from the confusion matrix (Figure 3e), the poor performance of the Sparrowhawk classification is predominantly responsible for lowering the precision and recall of the other species. Without the Sparrowhawk, the overall accuracy is significantly higher (0.5583 vs 0.7354: see Appendix B for Figure B.1 and Table B.1).

**Figure 3:**
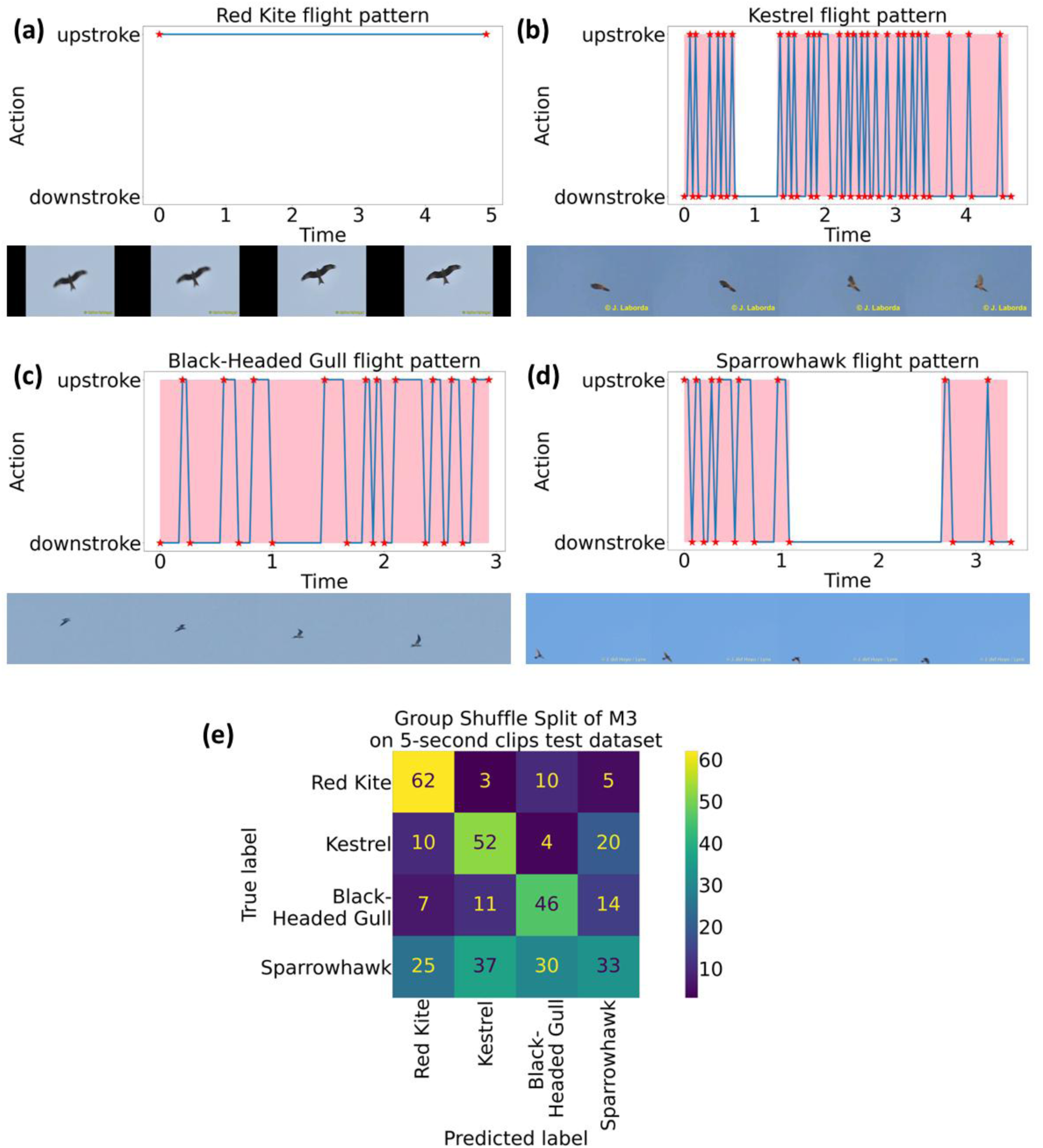
Feature extraction and species classification. (a)-(d) Examples of flight patterns with four frames from each clip used to generate each flight pattern showing the flapping or gliding of the bird. The switching points between upstrokes and downstrokes are highlighted by red star symbols and the identified flapping sections are highlighted by a pink background. (e) Confusion matrix for Group Shuffle Split cross-validation of M3 on the 5-second clips balanced dataset.

**Table 1:** Metrics per species for the Group Shuffle Split cross-validation of M3 on the 5-second clips balanced dataset.

| **Metric** | **Value** |  |  |  |
| --- | --- | --- | --- | --- |
| **Per Species** | Red Kite | Kestrel | Black-Headed Gull | Sparrowhawk |
| **Recall** | 0.7750 | 0.6047 | 0.5897 | 0.2640 |
| **Precision** | 0.5962 | 0.5045 | 0.5111 | 0.4583 |
| **F1 score** | 0.6739 | 0.5503 | 0.5476 | 0.3350 |

## 4. Discussion

The action classification model, M2, performed well, and upon further inspection many frames that were incorrectly classified as an upstroke when they were actually a downstroke (and vice versa) were at the switching points where the difference between actions is ambiguous. Figure 2f shows examples of a correctly classified upstroke and downstroke for each species in the 5-second clips test subset. The classification follows actual bird flight closely, with the upstroke frames showing the wings bent at the wing joint, and the downstroke frames showing the wings outstretched, despite the relatively poor resolution and focus of the input data. However, M2 performed even better when classifying the dataset of images in which the flying bird takes up most of the frame. M2 was trained on images more similar to those in the full-frame dataset, as the large-scale image datasets available for research contained images of that nature. Having training data that is more representative of real video data would allow our models to learn the correct features that apply to the actual application. For example, retraining M1 and M2 on a dataset comprising of images where the bird only takes up a small portion of the frame, like in the Zi-Wei Sun *et al* (2025) study (38) that comprises of 28,694 frames from surveillance video, might encourage the models to learn the features of the shape of the bird rather than the colour and appearance. This would be especially useful for M2 but would also allow for M1 to be more generalisable across more species and more contexts. Introducing data augmentation to the training datasets used for M1 and M2 would also be beneficial in improving their generalisability and preventing model overfitting, for example, by scaling, flipping, and rotating images to create pseudo-replicates, or by adding blur and modifying the brightness to simulate varying camera qualities and lighting conditions.

While the quality of training data is important, increasing the size of training datasets would also be beneficial, especially for the species identification model. The flight pattern types we chose for the proof-of-concept implementation usually belong to mid-size to larger birds, which have more video data available in crowd-sourced datasets than smaller birds because they are easier to identify from afar and capture on video.

The data was still quite limited, however, which has affected the performance of the model. This is most clearly seen in the poor recall of the Sparrowhawk, which had significantly fewer videos available in the original video dataset from the Macaulay Library (only 7 compared to the 40 available for the Red Kite), most of which were extremely similar or even derived from the same longer video, thereby resulting in significant overfitting during cross-validation, and under-representation of the natural differences in flight patterns between individual birds. Therefore, with a larger video dataset, it is possible that the performance would improve. However, the varying performance is also likely down to the performance of the previous stages. Subtle differences in the actual flight patterns of the Kestrel and the Black-Headed Gull, for instance, which come down to the timing of individual upstrokes and downstrokes, are likely being missed as a result of imperfect detection and classification of flight action. With more development to reduce these compounding errors, performance should improve and be more consistent across the species.

One major limitation of the current prototype system that limits its use in live applications is its processing time. Each 5-second clip takes approximately 4 minutes to process and obtain the species, using a high-end laptop with an Intel® Core™ i9-13900H 14 Core-Processor CPU, NVIDIA® GeForce RTX™ 3050 4GB GPU, and 16GB LPDDR5 RAM (39). To be deployed at scale, either on low-cost servers or on the edge (e.g., on single-board computers or microcontrollers within sensor devices), improving the computational efficiency is crucial. Processing time could potentially be reduced by parallelisation, but developing less complex computer vision models will likely make the largest impact, as the processing speed of M1 (and M2 to a lesser extent) is currently the main bottleneck of the system. Techniques for model compression have become an important research area in recent years. The main techniques are pruning, decomposition, quantization and distillation (40). Model compression for live bird detection appears to have not been demonstrated yet, as live bird detection is more commonly achieved using image detection which does not require as high of a processing speed since the capture rate of images is lower than the frame rate of videos. For example, the IdentiFlight system used in a study by McClure *et al* (2018) (41) captures images every 200ms, which is a capture rate of 5 images per second, whereas most of the videos we use in our approach have a frame rate of 25 fps, and therefore we need a processing speed that is at least five times faster than in an approach that solely uses appearance from images to identify species. However, one study by Rosmalen, Y (2025) (42) demonstrated the potential of model compression techniques in reducing the size of models by 98% for live automated detection of bears from camera traps. Future development of our approach could involve using model compression techniques to reduce the size of our bird detection (M1) and action classification (M2) models and therefore reduce processing time to the point that our approach can be used in live applications.

Despite the limitations of the current flight pattern generation process, the flight patterns generated in our proof-of-concept results do align with the actual types of flight patterns expected from the species chosen. Different species require different ratios of the flight modes (23) and of upstrokes to downstrokes (24) to produce the lift and thrust required for sustainable flight. This is a result of their biological traits, especially their size (43) and wing shape (44). Larger and heavier birds require more energy to produce lift when flapping than smaller, lighter birds, and therefore will glide more often (43). Different wing shapes are also better adapted to flapping or gliding flight (44). For example, hawks have large, broad wings to produce greater lift in updrafts, whereas ducks have long, pointed wings to improve the aerodynamics of continuous flapping motion (44). In Figure 3, we present an example of a generated flight pattern for each species. The Red Kite is a large bird-of-prey with broad wings, and this is clear from the generated flight pattern (Figure 3a), which shows gliding behaviour. Furthermore, in the original video, there was a lot of wind noise, possibly explaining why the pattern shows that the bird was flying in an upstroke glide rather than a downstroke, as this would have allowed for better control of lift to drag ratio in the face of strong winds. The Kestrel, on the other hand, is a smaller bird-of-prey that exhibits hovering behaviour when hunting. The flight pattern in Figure 3b shows this rapid flapping behaviour, with each upstroke and downstroke only taking around a fifteenth of a second. The more pointed wings of the Black-Headed Gull are designed for comfortable prolonged flapping motion but, with a larger wingspan than Kestrels, they can produce more lift and thrust with each downstroke. The flapping motion of the Black-Headed Gull is therefore slower, as seen in Figure 3c, with each upstroke and downstroke taking around a sixth of a second. Finally, the Sparrowhawk exhibits both flapping and gliding behaviour. With short but wide wings, it is capable of gliding for short periods but then requires rapid flapping motion to maintain its height and speed. This can be seen in the example flight pattern in Figure 3d.

Our approach has the potential to be used in many different applications, especially in combination with other detection and classification techniques to improve the overall accuracy of a species monitoring system. For example, to reduce the costs and required performance of visual equipment in monitoring systems that prevent bird strikes on wind farms, our method can be used to determine species from flight pattern where, unlike in turbine monitoring systems that use state-of-the-art camera systems to classify birds by appearance (41), the bird only takes up about 25 pixels and therefore its shape is the most distinguishable aspect of it (18). Beyond species identification, our approach to flight pattern extraction could be used by behavioural ecologists studying flight patterns in wild or captive birds to collect large-scale and longitudinal data. The health of protected birds under surveillance (45) could also potentially be monitored as changes in their flight pattern may indicate injury (46).

## 5. Conclusion

In this study, we presented a novel approach to characterising the flight patterns of birds by analysing easily and inexpensively acquired videos from non-specialist cameras with a combination of off-the-shelf and custom machine learning models. Beyond just species identification, our approach can also output important data on bird flight that can be used to study the behaviour and monitor the health of individuals in the wild and in captivity. Although we have demonstrated a small-scale proof-of-concept system, we do anticipate that with higher fidelity datasets covering a broader set of species, our novel approach will be able to provide a useful additional dimension to bird monitoring efforts across a wide range of real-world applications.

## Data Availability Statement

The project code has been made available as a release on Zenodo via https://doi.org/10.5281/zenodo.18332340. The labels for the M2 training dataset have been made available on Zenodo via https://doi.org/10.5281/zenodo.18281931. The images the labels correspond to can be downloaded from the original datasets: NABirds (https://dl.allaboutbirds.org/nabirds) and iNaturalist 2019 (https://www.kaggle.com/competitions/inaturalist-2019-fgvc6/data). The labels for the M2 full-frame test dataset have been made available on Zenodo at https://doi.org/10.5281/zenodo.18282125. The images the labels correspond to can be downloaded from the original dataset: CUB-200-2011 (https://www.vision.caltech.edu/datasets/cub_200_2011/). The other flying objects dataset has been made available on Zenodo via https://doi.org/10.5281/zenodo.18281801. We also used videos from the Macaulay Library that we are not able to share, but a list of the catalogue numbers is available in the Appendix so that the same videos can be requested.

## Acknowledgments

We thank the Cornell Lab of Ornithology for providing access to videos from the Macaulay Library and for providing access to the NABirds dataset. We thank the Caltech Vision Lab (Perona Lab) at the California Institute of Technology for providing access to the CUB-200-2011 dataset. We thank iNaturalist and FGVC6 for providing access to the iNaturalist 2019 dataset.

## Declaration of generative AI and AI-assisted technologies in the manuscript preparation process

No generative AI or AI-assisted technologies were used in the manuscript preparation process.

## Appendix A. Object Tracking

**Figure A.1:**
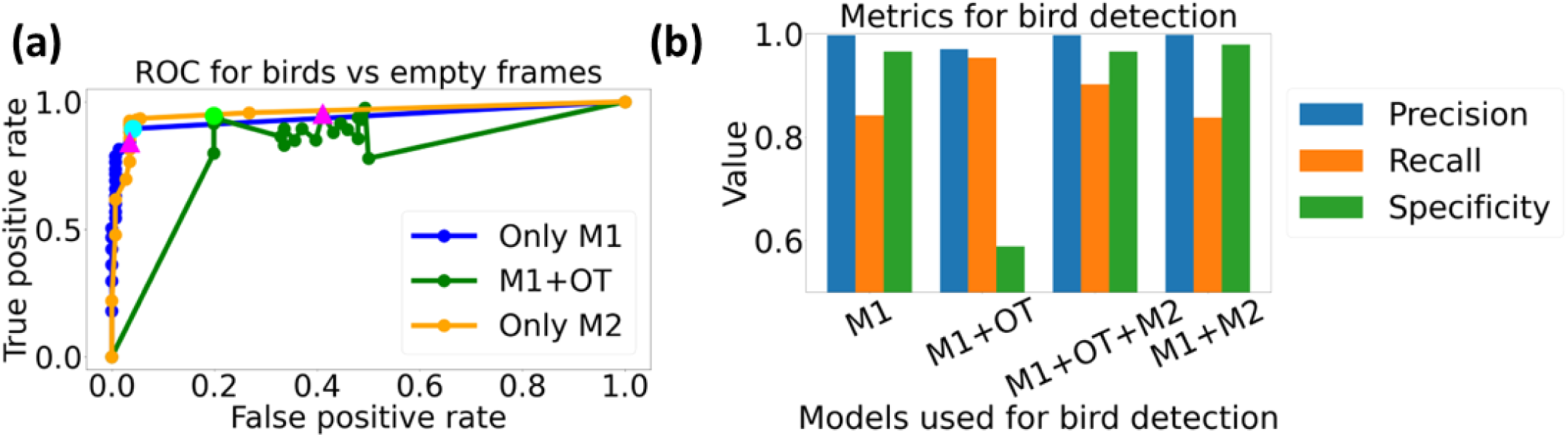
(a) Receiver Operating Characteristic (ROC) for the 5-second clips test subset, classifying birds vs empty frames, for M1 only (blue), M1+OT (green) and M2 only (orange) with the optimal thresholds highlighted by cyan (M1) and lime (M1+OT) circles (Only M1: 0.05, M1+OT: 0.15). We use M1 at a threshold of 0.1 to utilise its ability to identify other flying objects. A threshold of 0.1 is highlighted on the M1 (blue) and M1+OT (green) lines with magenta triangles. A threshold of 0.1 for M1 results in a drop in true positive rate compared to a threshold of 0.05, and for M1+OT, a threshold of 0.1 results in a significant increase in false positive rate. Unlike for M1 only and M2 only, the points on the ROC for M1+OT are not in order of ascending threshold going right to left. This suggests that the performance of object tracking is heavily dependent on the specific combination of frames that M1 designates as a bird, making it unpredictable for new data. (b) Bar chart showing precision, recall and specificity of each stage of bird detection. For solely M1 used at a threshold of 0.1, precision is 0.9971, recall is 0.8425, and specificity is 0.9658. For M1 + OT, precision is 0.9706, recall is 0.9538 and specificity is 0.5890. For M1 + OT + M2, precision is 0.9973, recall is 0.9022 and specificity is 0.9658. For M1 + M2, precision is 0.9983, recall is 0.8382, and specificity is 0.9795. M1 + OT + M2 has a higher recall but slightly lower specificity than M1 + M2. As there isn’t a hugely significant difference between performance when including or excluding OT, and OT seems to be heavily dependent on the specific frames involved, we opt to just use M1 + M2.

## Appendix B. No Sparrowhawk

**Figure B.1:**
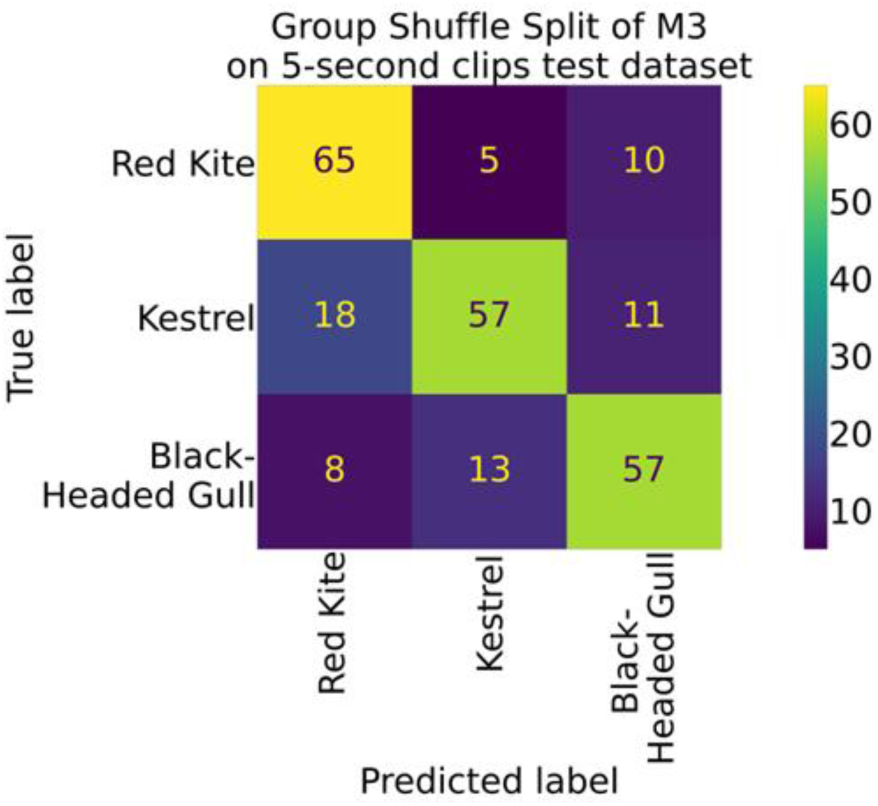
Confusion matrix for Group Shuffle Split cross-validation of M3 on the 5-second clips balanced dataset without the Sparrowhawk.

**Table B.1:**
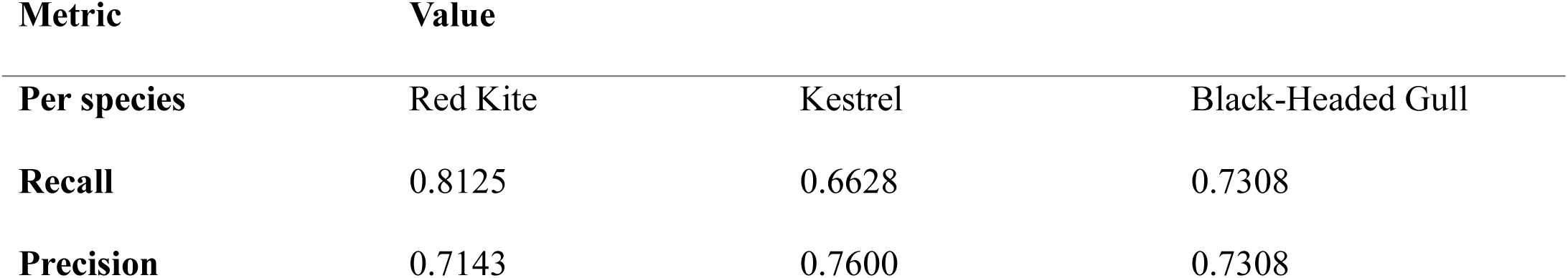
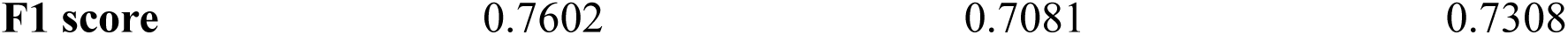
Table showing the metrics for the Group Shuffle Split cross-validation of M3 on the 5-second clips balanced dataset without the Sparrowhawk. All the metrics are significantly higher than when the Sparrowhawk is included, indicating that the lack of variation in the Sparrowhawk dataset is the most detrimental factor to the performance of species identification.

## Appendix C: Macaulay Library Catalogue Number list

I used the following recordings from the Macaulay Library at the Cornell Lab of Ornithology: 201078921, 201084431, 201421621, 201435811, 201466721, 201471701, 201471711, 201475381, 201489841, 201495771, 201496841, 201510741, 201510751, 201512931, 201530571, 201734091, 201766941, 201815221, 201857941, 201882221, 201898741, 201907461, 201924041, 201926521, 201926571, 217728711, 223173511, 242197251, 323863541, 460397631, 466608321, 201233861, 201387831, 201466901, 201492201, 201520471, 201852131, 201873771, 201907511, 201907531, 201907551, 201907571, 201909401, 201931481, 247632411, 276429291, 388243571, 513057611, 413776, 201411291, 201496651, 201496661, 201853641, 201853641, 201864421, 201487651, 201487661, 201490041, 201499601.

Some videos contained multiple clips with transitions, and so were manually split.

